# MAUDE-Py: A Python Implementation for Analyzing Sorting-Based Pooled CRISPR Screens

**DOI:** 10.64898/2026.11.12.766045

**Authors:** Avirral Agarwal, Jennifer L. Moy

## Abstract

The Python programming language is gaining popularity in bioinformatics owing to its versatility in computing-heavy applications. An area of particular interest where computational power can be leveraged is analyzing and interpreting pooled CRISPR screens. Here, we replicate the core functionality of an existing platform, Mean Alterations Using Discrete Expression (MAUDE), in Python (MAUDE-Py) to analyze pooled CRISPR screens. Our tool complements the original R MAUDE software as a robust and user-friendly Python package that integrates with existing CRISPR analysis workflows. While preserving the core probabilistic framework of MAUDE, MAUDE-Py introduces notable enhancements. These enhancements include automated baseline calculations, a more robust null-sampling methodology for determining significance at both the gene and non-coding element levels, and an integrated visualization suite for displaying the analyzed results. Benchmarking MAUDE-Py against the original MAUDE on a series of datasets demonstrates a 2.09× to 4.19× speedup in execution time while achieving near-perfect consistency, confirming that our Python implementation is a highly accurate and viable alternative for such analyses.

## Introduction

Clustered Regularly Interspaced Short Palindromic Repeats (CRISPR) screens have emerged as a foundational technology in functional genomics, largely replacing traditional methods such as RNA interference (RNAi) due to their improved specificity, scalability, and efficiency [1]. In arrayed CRISPR screens, individual single guide RNAs (sgRNAs or gRNAs) are placed into separate wells to knock out, knock down, or activate target genes [2–4]. Because scaling this setup is expensive and labor-intensive, pooled CRISPR screens have been developed to bypass these bottlenecks by delivering an entire gRNA library into a single population of cells [5]. However, because the cells are mixed together, linking individual gRNAs to their specific phenotypic shifts requires downstream computational analysis.

To measure specific cellular responses in pooled formats, cells are engineered to express a fluorescent reporter, such as Green Fluorescent Protein (GFP), that is driven by a pathway of interest before being treated with a gRNA library and separated into discrete expression bins via Fluorescence-Activated Cell Sorting (FACS) [5–7]. Following sorting, the gRNAs within each bin are amplified and sequenced to generate guide count tables across the expression distribution. While CRISPR analysis tools such as Model-based Analysis of Genome-wide CRISPR-Cas9 Knockout (MAGeCK) [8], PinAPL-Py [9], and ScreenPro2 [10] exist, these are primarily for selection-based screens. They may, for example, be used for analyzing enrichment and depletion data between discrete conditions, such as treatment versus control data. However, such tools are not meant for sorting-based screens, which involve continuous phenotypic data like fluorescence expression across bin ranges derived from FACS. Mean Alterations Using Discrete Expression (MAUDE) is a tool designed specifically for analyzing such CRISPR screening data [11].

While MAUDE can provide robust and comprehensive analyses for sorting-based screens, its implementation in the R programming language presents a barrier for those with limited exposure to the language or who are not using R in their current workflow. As an alternative to R, the Python programming language is gaining traction in bioinformatics and computing due to its simple syntax and rich ecosystem support [12, 13]. This motivates our development of MAUDE-Py, a Python version of MAUDE that replicates the tool’s core behavior of analyzing sorting-based pooled CRISPR screen data. MAUDE-Py is designed to complement MAUDE by supporting existing workflows in Python while maintaining its core logic, with some added functionality compared to the native R version to improve usability.

## Methods

### Statistical Framework and Algorithmic Implementation

MAUDE-Py implements fundamental statistical concepts and models to identify significant changes in the cells’ phenotype, as measured by GFP expression, replicating the behavior of the original MAUDE [11]. The program first processes input tables containing gRNA counts across GFP expression bins in addition to corresponding target identifiers, such as genes or non-coding genomic positional data. User-defined percentile ranges for GFP expression bins that are based on experimental FACS conditions are converted into Z-scores for use in the core probabilistic predictive model. Similar to the original implementation, MAUDE-Py includes the “makeBinModel” function to convert the fraction of cells captured in each bin into cumulative percentile ranges when not provided directly. Additionally, MAUDE-Py includes error handling for missing experimental data. If an unsorted baseline read column is not provided, MAUDE-Py generates one by averaging the counts across all bins for that gRNA and rounding to the nearest integer. This feature is not present in the original MAUDE.

Before a model can be fit to estimate the effects of the gRNAs, we need to account for natural variability in the data. The Poisson model is often proposed to estimate such processes due to its simplicity in assuming the variance equals the mean; however, this model fails to account for the fact that next-generation sequencing (NGS) screening data often has a sample variance greater than the mean [8]. The negative binomial model accounts for this difference and is standard in CRISPR and RNA-seq analysis tools such as MAGeCK [8], edgeR [14], and DESeq [15].

As discussed by Li et al. [8], the negative binomial dispersion parameter, *k*, models data variability and can be estimated using the method of moments approach. For a negative binomial distribution, parameterized as *NB*(*k, p*), the mean is 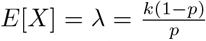 and the variance is 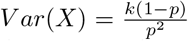[16]. By isolating *p* and substituting it into the variance equation, the variance can be re-expressed in terms of the mean and the dispersion parameter as

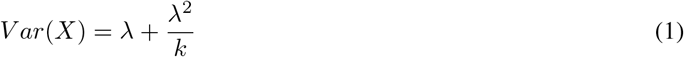

The original MAUDE implementation uses the negative binomial model, defining the dispersion parameter *k* as the total counts in the GFP expression bins [11]. Because any single guide makes up only a tiny fraction of the total reads in a bin, a guide’s expected count *λ* is several orders of magnitude smaller than the total bin count *k* (*λ* ≪*k*), making the ^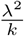^ term negligible compared to *λ*. Thus, we can make the approximation that *V ar*(*X*) = *λ*. This is the defining property of the Poisson distribution [16]. Using the Poisson distribution is appropriate here because sorting-based screens compare cell fractions within a single experiment rather than comparing read counts across different samples, naturally avoiding the extreme overdispersion commonly seen in standard RNA-seq data. Although it is possible that different bins may be sequenced at different depths, this is already accounted for in downstream processing. Based on these ideas, MAUDE-Py skips the negative binomial calculation entirely and instead directly uses the Poisson distribution for modeling the data.

The core logic of MAUDE is a probabilistic model based on the standard normal distribution to quantify the phenotypic effect of each gRNA [11]. The goal is to find a value, *shift µ*, for each guide that causes the expected counts — the model’s predicted reads for a guide — to match the guide’s observed counts for each bin. First, the model calculates the expected fraction of reads in bin *b* for guide *g*, denoted as *P*_*b*_(*g*| *µ*_*g*_), which matches equation (2) in the original MAUDE paper [11]:

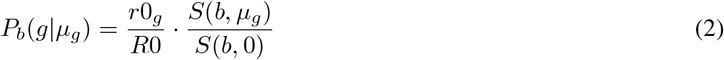

Here, *r*0_*g*_*/R*0 accounts for sequencing depth and represents the proportion of unsorted input (“background”) reads relative to the total number of unsorted input reads. *S*(*b, µ*_*g*_) and *S*(*b*, 0) are computations for probabilities under the standard normal distribution, with *S*(*b, µ*_*g*_) representing the fraction of data that is expected to lie in the bin with a given *shift µ* and *S*(*b*, 0) representing the fraction of data expected to lie in the bin with mean 0 and standard deviation 1. Their ratio represents the fraction of data that would lie in the bin given a particular *shift µ* when compared to the baseline null curve. Multiplying the *P*_*b*_(*g*| *µ*_*g*_) term by *R*_*b*_, the total number of reads in a bin, gives a value representing the expected number of reads for guide *g* in each bin given a particular *shift µ*.

Finding expected counts allows MAUDE-Py to compare against the actual observed counts from experiments through the log-likelihood function. Log-likelihood wraps the probability distribution in a logarithmic formula, as multiplying small probabilities for each bin leads to underflow [17]. We maximize the log-likelihood function through repeated cycles of refining *shift µ* to gain a better approximation of expected counts that more closely matches the observed counts. As stated in the MAUDE paper, the original platform maximizes the log-likelihood using the negative binomial distribution [11]. Because we established that the Poisson distribution appropriately models the data for sorting-based screens, MAUDE-Py instead maximizes the following equation for each guide:

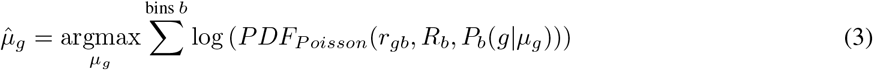

For computational efficiency, we minimize the negative log-likelihood, which is mathematically the same as maximizing the log-likelihood. We identify the best *shift µ* for each guide using SciPy’s bounded scalar minimizer (Brent’s method) [18], which differs from the original MAUDE’s gradient-based L-BFGS-B method [11]. By subtracting the mean of the non-targeting control (NTC) guides from each guide’s optimized *shift µ*, these values are converted to normalized Z-scores, as modeled by equation (4) of the MAUDE paper [11].

At the gene level, we quantify the combined effect of all associated gRNAs for the same gene by calculating a Stouffer Z value, in accordance with Stouffer’s formula [19]:

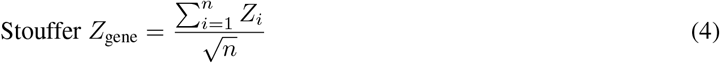

Here, *n* represents the number of guides targeting the gene and *Z*_*i*_ represents the normalized Z-score of each guide. To evaluate gene significance, we apply an empirical null model to establish a baseline that accounts for variability among NTC guide Z-scores [11]. While it is expected that NTC guides have a mean of 0 and a standard deviation of 1, experimental noise may cause slight variations from these values in the observed data. For each gene size, defined as the number of gRNAs associated with a gene, MAUDE-Py simulates an experiment where NTC guides are sampled at random with replacement. Stouffer Z-scores are calculated for these sampled groups, and their standard deviation serves as the null standard deviation (*σ*_0_). Because guide-level scores were already centered around the NTC mean during initial optimization, the expected null mean is zero. We then compute the final “significanceZ” for each gene by dividing the gene Z-score by *σ*_0_:

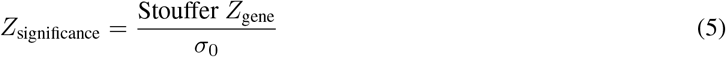

These “significanceZ” scores represent, in the form of a Z-score, the number of standard deviations the guides from a particular gene shift expression compared to random noise. To account for multiple hypothesis testing across thousands of genes, MAUDE-Py converts these “significanceZ” scores into two-tailed p-values and uses the Benjamini-Hochberg method [20] to compute a false discovery rate (FDR) for each gene.

The sampling approach MAUDE-Py uses offers an improvement over the original MAUDE implementation. The original MAUDE creates a fixed pool of NTC guides by multiplying the total number of controls by a constant before shuffling that pool and slicing it by gene size to generate null groups. Because it slices a fixed pool without replacement, the number of simulated null groups decreases as gene size increases. By sampling with replacement instead, MAUDE-Py ensures that larger gene sizes maintain sufficient simulated null groups, yielding more consistent estimations.

Similar to the original MAUDE, we also define methods to process CRISPR data for experiments that target non-coding regions of the genome, such as enhancers and promoters. Two such methods are implemented: an element-wise method that evaluates guides mapping to predefined genomic windows, and a sliding-window method that aggregates overlapping guides across contiguous intervals. The sliding-window approach is designed to group adjacent guides together to process them as pseudo-genes, outputting significance metrics consistent with gene-level statistics. Additionally, MAUDE-Py supports multi-experiment handling by analyzing experimental replicates independently so that inter-replicate variability does not confound the final analysis. All these methods are analogous to the original MAUDE in design and logic.

### Visualizations

To aid in interpreting the results, we developed a visualization suite. The visualization suite is designed to work only with CRISPR screens that target coding regions of the genome. After running the core CRISPR screen analysis, we extract the guide-level effects (“Z-score”, “Gene”, and “sgRNA”) and gene-level summary metrics (“meanZ” and “significanceZ”). To confirm that the “significanceZ” center around zero and approximate a standard normal distribution, which is necessary for downstream statistical analysis, we plot a density curve of the gene-level “significanceZ” scores.

In addition, we highlight the genes exhibiting the strongest biological responses. We identify the genes whose guides exhibit the largest average shifts in expression, considering both positive and negative shifts. We plot individual gRNA Z-scores for these top genes, and also generate a box-and-whisker plot depicting the 25th, 50th (median), and 75th percentiles of the guide-level Z-scores for each gene. We also overlay the box-and-whisker plots with the mean of the Z-scores (“meanZ”) for each respective gene.

We generate a similar plot for the genes that have the least change in biological response; this plot displays the genes with a mean guide-level expression shift (“meanZ”) closest to zero.

### Benchmarking

To evaluate the performance of MAUDE-Py, we included methods to compare the results obtained by MAUDE-Py with those obtained from the original MAUDE. We then computed statistical metrics to quantify differences between the results, as discussed below.

### Generative AI Statement

During the preparation of this work, the authors used Generative AI models by OpenAI and Google Gemini to assist with drafting the initial Python code and benchmarking scripts. Following this, the authors manually verified all mathematical formulas, designed the object-oriented architecture, optimized the code, and wrote the manuscript. The authors take full responsibility for the content, logic, and accuracy of the final software and publication.

## Results

For the purposes of comparison and benchmarking, we used both the original MAUDE and MAUDE-Py to analyze four CRISPR screens: simulated data generated via the original MAUDE simulation tutorial [11], CD69 CRISPRa data [21], BACH2 base editor data [22], and genome-wide LOUP CRISPRi screen data (GEO accession: GSE247761) [23]. The first three of these data sources were discussed in tutorials for the original MAUDE [11], ensuring the R code used for preprocessing was consistent with the original framework. Preprocessing of the raw data for all screens was performed in R, and the identically processed data was provided as input to both MAUDE and MAUDE-Py to ensure consistency.

To quantify the alignment between the two implementations, we computed the Pearson correlation, Spearman correlation, and the Root Mean Square Error (RMSE). The Pearson correlation measures the strength and direction of the linear relationship between the two variables, ranging from -1 to +1 [24, 25]. RMSE measures the square root of the mean of the squared residuals between the values of the datasets [26], while the Spearman correlation ranges from -1 to +1 and measures relative rank ordering [27, 28].

Across all four sets of data, we observed high degrees of alignment in outputs. The comparative analysis yielded Pearson coefficients ranging from 0.9871 to 1.0000, Spearman coefficients ranging from 0.9793 to 1.0000, and RMSE ranging from 0.0000 to 0.3132 (Figs. 1–4). All datasets were analyzed at both the individual guide and gene or genomic element levels, excluding the BACH2 dataset. The BACH2 dataset was intentionally evaluated only at the guide level without gene-level aggregation, since base-editing screens evaluate the functional impact of precise, individual point mutations against a wild-type baseline.

**Figure 1:**
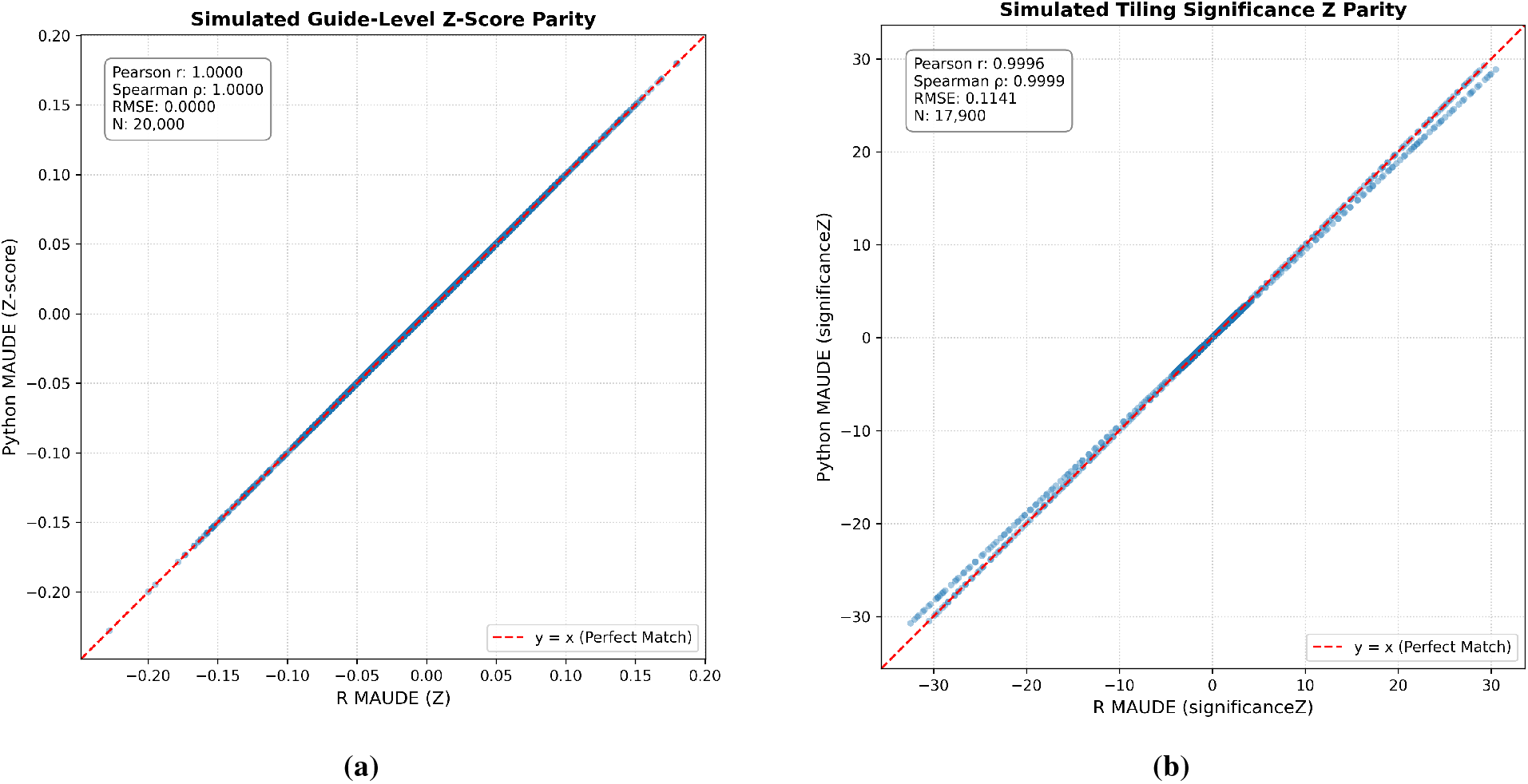
Scatter plots comparing the original MAUDE and MAUDE-Py performance on the Simulated Dataset. **(a)** The scatter plot for Z-scores at the guide level shows perfect alignment between MAUDE and MAUDE-Py, with a Pearson coefficient of 1.0000, a Spearman coefficient of 1.0000, and an RMSE of 0.0000. **(b)** The scatter plot for “significanceZ” scores using the tiling (sliding-window) method shows near-perfect alignment between MAUDE and MAUDE-Py, with a Pearson coefficient of 0.9996, a Spearman coefficient of 0.9999, and an RMSE of 0.1141.

**Figure 2:**
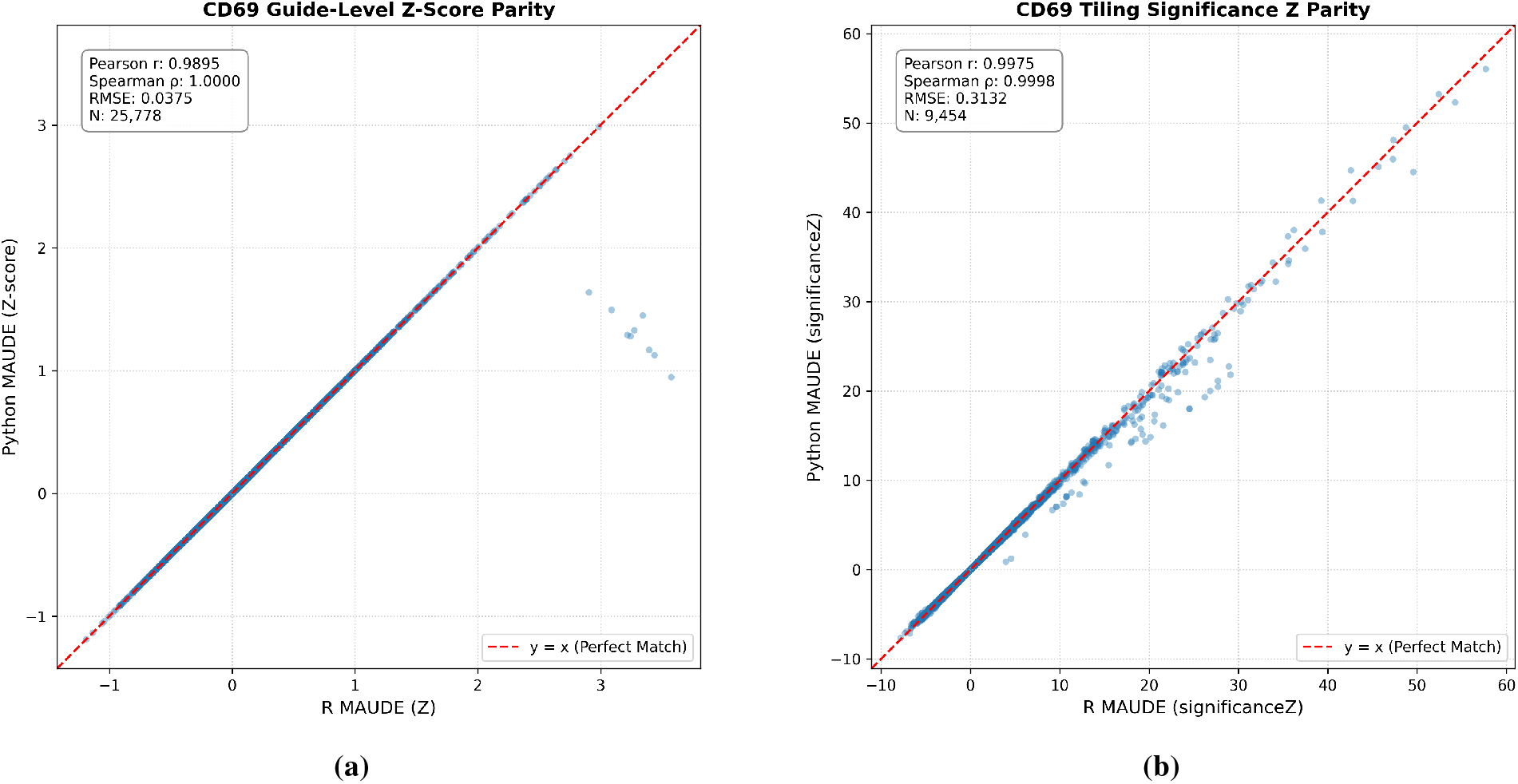
Scatter plots comparing the original MAUDE and MAUDE-Py performance on CD69 CRISPRa Data. **(a)**The scatter plot for Z-scores at the guide level shows near-perfect alignment between MAUDE and MAUDE-Py, with a Pearson coefficient of 0.9895, a Spearman coefficient of 1.0000, and an RMSE of 0.0375. **(b)** The scatter plot for “significanceZ” scores using the tiling (sliding-window) method shows near-perfect alignment between MAUDE and MAUDE-Py, with a Pearson coefficient of 0.9975, a Spearman coefficient of 0.9998, and an RMSE of 0.3132.

**Figure 3:**
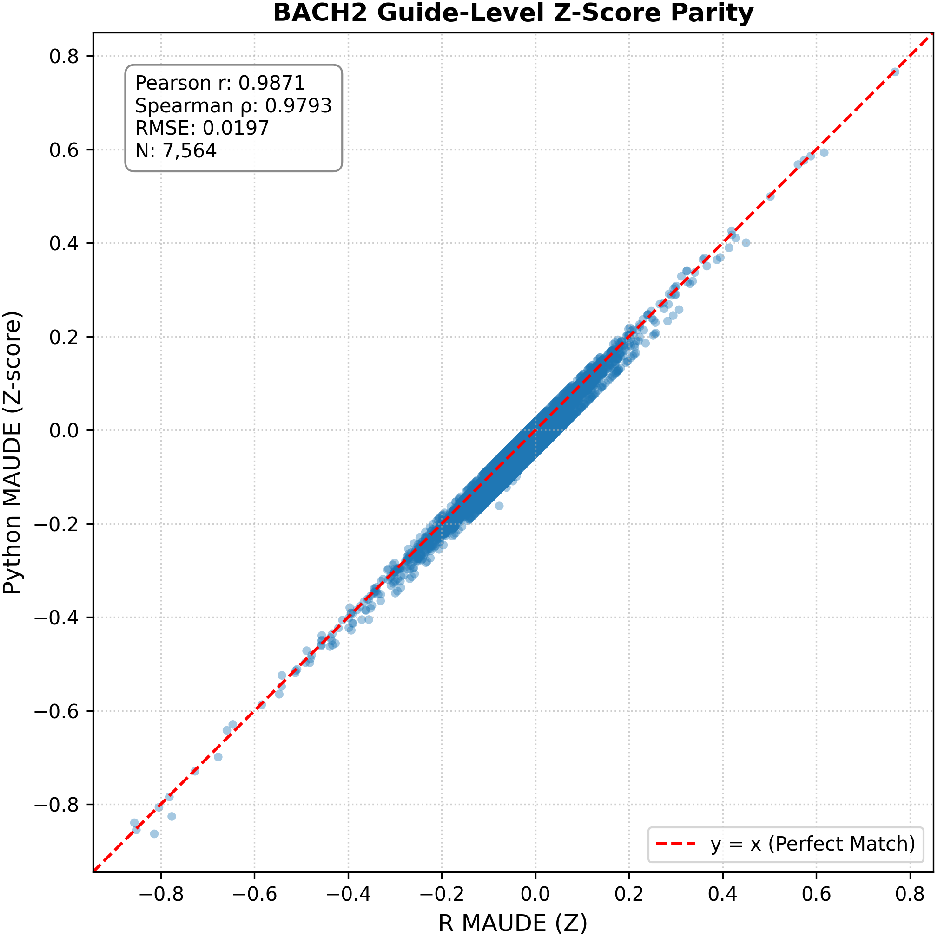
Scatter plot comparing the original MAUDE and MAUDE-Py performance on the BACH2 Dataset. The scatter plot for Z-scores at the guide level shows near-perfect alignment between MAUDE and MAUDE-Py, with a Pearson coefficient of 0.9871, a Spearman coefficient of 0.9793, and an RMSE of 0.0197. Gene-level aggregations are omitted for this dataset, as individual base-editing variants must be evaluated independently against their wild-type baseline.

**Figure 4:**
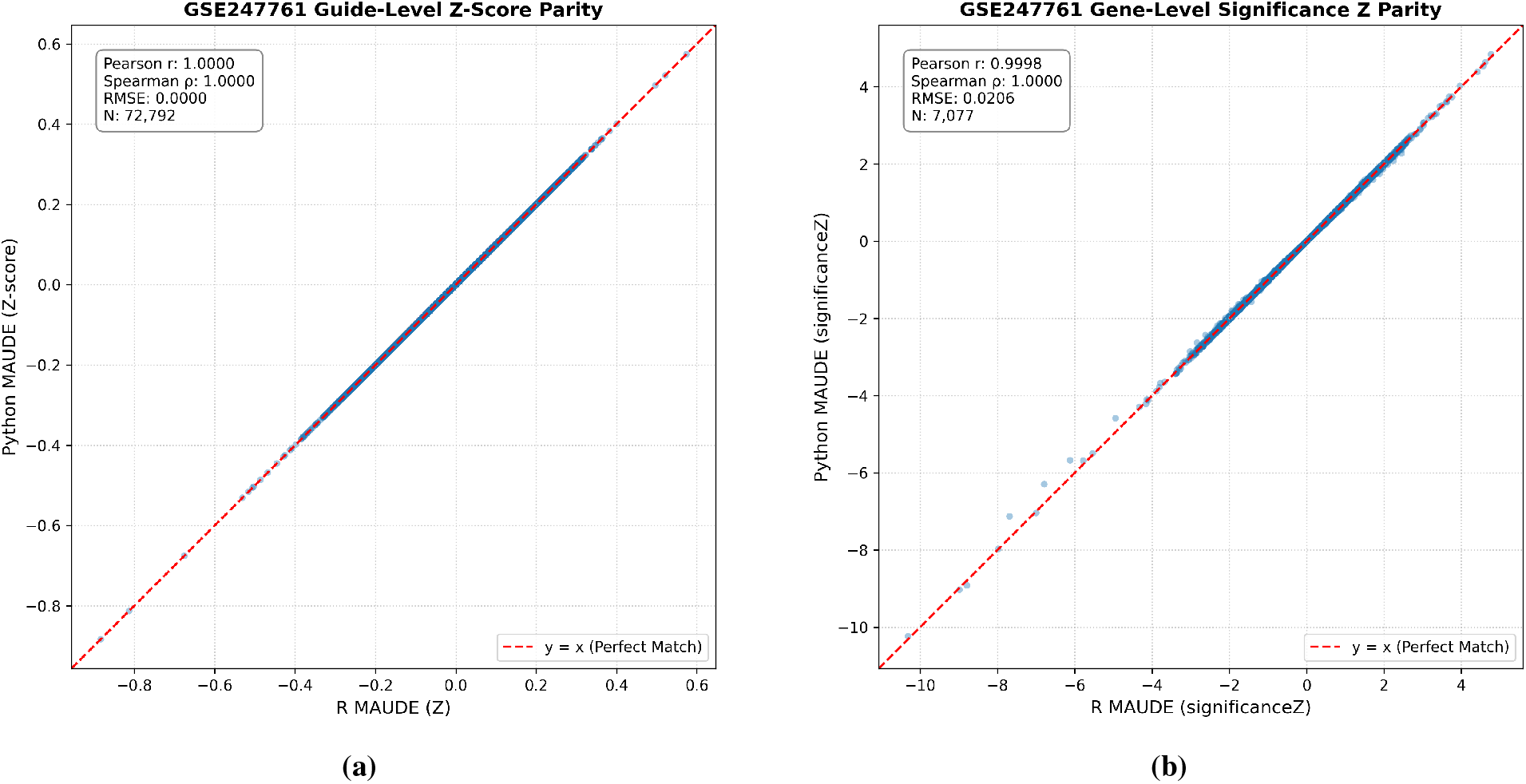
Scatter plots comparing the original MAUDE and MAUDE-Py performance on the CRISPRi dataset. **(a)** The scatter plot for Z-scores at the guide level shows perfect alignment between MAUDE and MAUDE-Py, with a Pearson coefficient of 1.0000, a Spearman coefficient of 1.0000, and an RMSE of 0.0000. **(b)** The scatter plot for “significanceZ” scores at the gene level shows near-perfect alignment between MAUDE and MAUDE-Py, with a Pearson coefficient of 0.9998, a Spearman coefficient of 1.0000, and an RMSE of 0.0206.

**Figure 5:**
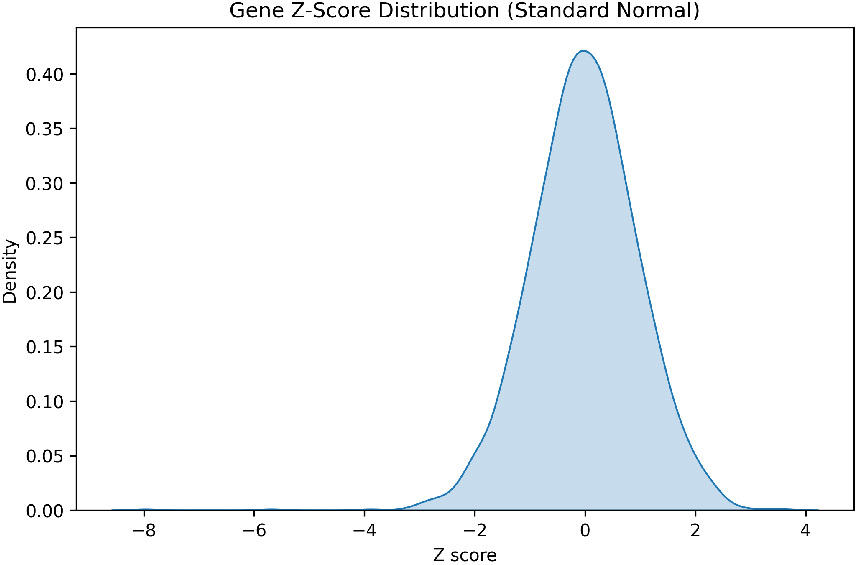
Density curve of the gene-level “significanceZ” scores. The distribution resembles a standard normal curve centered around 0, confirming the background data is correctly scaled for downstream p-value and FDR calculations. The second component of our visualization suite is a method to plot genes that caused the largest change in expression compared to the baseline. To highlight the most pronounced phenotypic shifts, the pipeline ranks genes by the absolute magnitude of their average guide-level shift (“| meanZ |”) and displays the top 20 genes with the largest overall expression changes (Fig. 6). The final component of our visualization suite is a method to plot the genes that caused the least change in expression compared to the baseline. We select the 20 genes with “| meanZ |” values closest to zero (Fig. 7), illustrating that these genes and their associated gRNAs center around zero.

**Figure 6:**
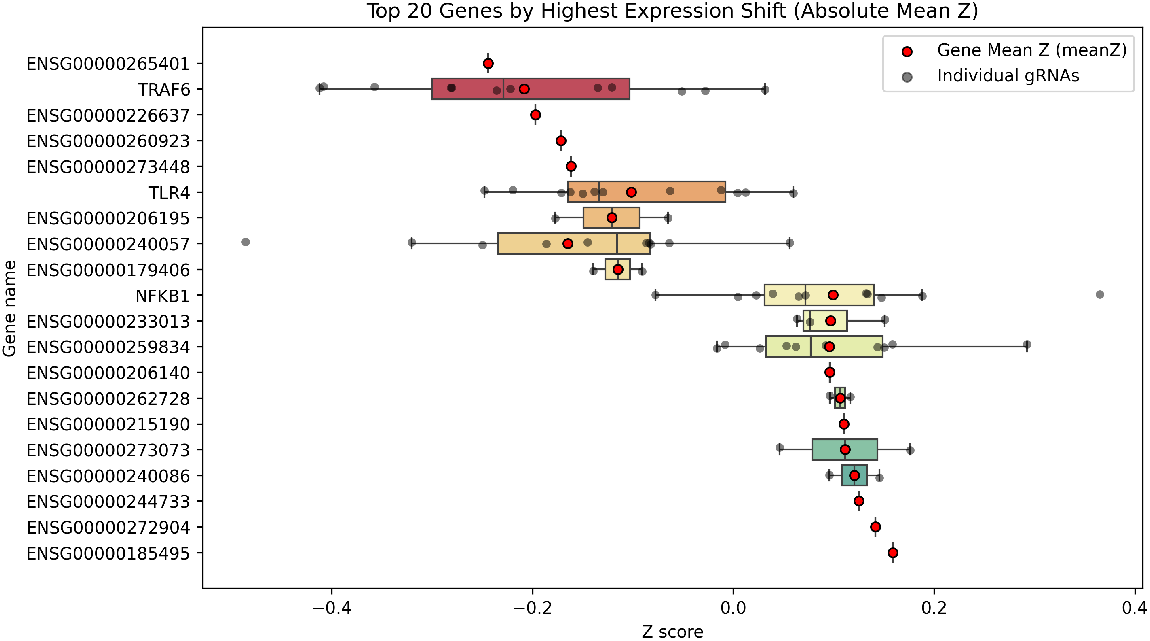
Plot of the genes with the largest average expression shift by absolute mean effect size. A box-and-whisker plot displaying the five-number summary for gRNA Z-scores associated with the 20 genes that had the highest mean expression shifts (regardless of whether they were positive or negative) for the CRISPRi screening data. The plot also includes the mean of the Z-scores for each gene. Genes targeted by a single gRNA appear as individual points without interquartile boxes.

**Figure 7:**
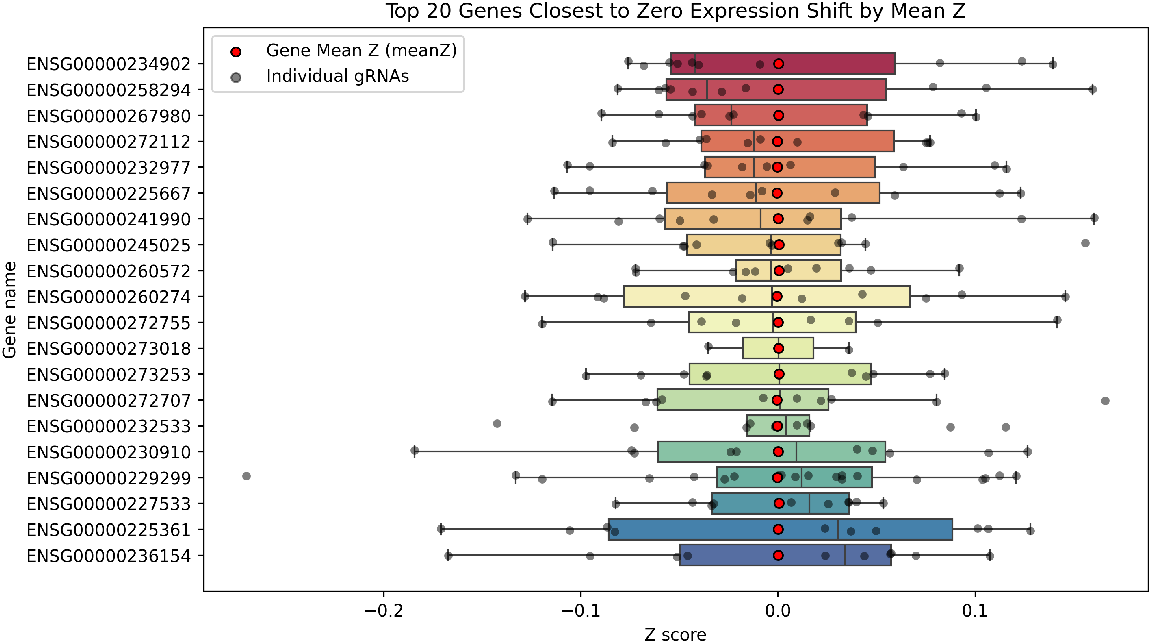
Plot of the genes with the average expression shift closest to zero. A box-and-whisker plot displaying the five-number summary for gRNA Z-scores associated with the 20 genes that had mean expression shifts closest to zero for the CRISPRi screening data. The plot also includes the mean of the Z-scores for each gene.

Furthermore, we recorded the execution time for the core analytical pipeline, excluding preprocessing and visualization, for both MAUDE and MAUDE-Py on identical hardware (Table 1). Across all four datasets, MAUDE-Py consistently outperformed the native R implementation, achieving a 2.09× to 4.19× speedup in execution time. This speedup is largely attributable to aspects such as using NumPy matrix operations for expected-count evaluations and empirical null sampling, using a different optimizer function, and using the Poisson distribution for computing log-likelihood.

**Table 1:** A comparison of computation time between MAUDE-Py and the original MAUDE on the four bench-marking datasets. MAUDE-Py and MAUDE were both run locally on a 2022 MacBook Pro equipped with an Apple M2 processor and 8 GB of RAM. Execution times represent the core analytical pipeline, excluding preprocessing and downstream plotting. Across all four datasets, MAUDE-Py demonstrates notably faster processing times than MAUDE. The first, a method to plot the density curve to assess normality of the data, is utilized to show the distribution of the “significanceZ” scores at the gene level. The distribution resembles a standard normal curve centered around zero (Fig. 5). This is essential, as genome-wide statistical significance is determined by p-values and FDR thresholds that utilize the standard normal distribution.

| **Dataset** | **Time for MAUDE-Py (s)** | **Time for MAUDE (s)** | **Speedup Factor<br>(MAUDE/MAUDE-Py)** |
|---|---|---|---|
| Simulated | 19.64 | 82.29 | 4.19× |
| CD69 | 27.69 | 100.03 | 3.61× |
| BACH2 | 7.34 | 16.15 | 2.20× |
| CRISPRi | 64.83 | 135.48 | 2.09× |

To demonstrate the use of MAUDE-Py’s visualization suite, we inspected the results of running MAUDE-Py on Replicate A of the CRISPRi screen dataset (Clone A) [23]. The visualization suite has three main components.

## Discussion

Pooled CRISPR screens have become central to understanding transcriptional regulation and non-coding genomics. However, widely adopted tools like the original MAUDE package have historically been confined to the R ecosystem. MAUDE-Py addresses this gap by providing an open-source, pure-Python implementation of the MAUDE framework, integrating seamlessly into modern Python data science environments such as Google Colab, Jupyter, and standard bioinformatics workflow managers.

Across multiple benchmark datasets encompassing both simulated and biological screens (including activation, base-editing, and interference assays), MAUDE-Py achieves outputs nearly identical to those of the original MAUDE. MAUDE-Py additionally introduces refinements to the underlying workflow. In the null estimation step, sampling NTCs with replacement resolves a limitation in the original package, where slicing a fixed-size control pool reduced the number of simulated null groups as gene sizes increased. MAUDE-Py maintains a uniform null simulation sample size across all gene sizes. Additionally, MAUDE-Py uses the Poisson distribution directly instead of MAUDE’s approach of parameterizing a negative binomial distribution that collapses to a Poisson distribution. In our benchmark comparisons across experimental datasets, this formulation yielded virtually indistinguishable effect estimates and correlation metrics.

Areas of additional work include improving the visualization suite and further improving computational speed and performance. In regard to the former, expanding the visualization suite to handle non-coding CRISPR screen data may prove useful. In addition, displaying only genes that are statistically significant, i.e. they pass a certain “significanceZ”, p-value, or FDR threshold, would likely improve the biological interpretability of the analysis. In terms of speed and performance, the optimization function to find the best *shift µ* that maximizes the log-likelihood can be improved. While we use an optimizer from the SciPy library [18], decreasing error tolerance or changing the optimizer entirely may further improve runtime performance with limited to no negative effect on result accuracy. Further opportunities to increase speed and performance could include adding parallelization and Graphics Processing Unit (GPU) support, as well as using the Numba Just-in-Time (JIT) compiler [29].

While we verified MAUDE-Py across four diverse screening modalities, including synthetic, CRISPRa tiling, base-editing, and genome-wide CRISPRi datasets, it would be worthwhile to verify accuracy by comparing performance against more screens — perhaps those that have more variation in FACS bins and other experimental conditions — to more accurately and comprehensively benchmark model performance.

## Conclusion

MAUDE-Py is a comprehensive adaptation of MAUDE in the Python programming language designed to analyze CRISPR screens that use FACS methods. MAUDE-Py employs the same core statistical processes and functionality used by the original MAUDE while introducing improved null sampling, missing-data handling, and visualization capabilities.

With interest and research in computing and bioinformatics growing, the Python programming language is becoming more widely utilized. Our hope is that MAUDE-Py will complement the existing MAUDE software and be adopted for pooled CRISPR screen analysis.

## List of Abbreviations

CRISPR: Clustered Regularly Interspaced Short Palindromic Repeats
FACS: Fluorescence-Activated Cell Sorting
FDR: False Discovery Rate
GFP: Green Fluorescent Protein
GPU: Graphics Processing Unit
gRNA: Guide RNA
JIT: Just-in-Time
MAGeCK: Model-based Analysis of Genome-wide CRISPR-Cas9 Knockout
MAUDE: Mean Alterations Using Discrete Expression
MAUDE-Py: Mean Alterations Using Discrete Expression in Python
NGS: Next Generation Sequencing
NTC: Non-Targeting Control
RMSE: Root Mean Square Error
RNAi: RNA interference
sgRNA: Single Guide RNA

## Declarations

### Ethics approval and consent to participate

Not applicable.

### Consent for publication

Not applicable

### Authors’ Contributions

AA developed the initial MAUDE-Py codebase, implemented the benchmarking script, and tested the benchmarking methods on the various datasets. Both AA and JM developed the visualization suite, conceived and implemented the new class design for MAUDE-Py, and drafted and edited the manuscript. All authors read and approved the final manuscript.

### Availability of data and materials

All data and code pertaining to MAUDE-Py, visualizations, and benchmarking are available in the GitHub repository: https://github.com/aviaga/MAUDE-Py.

